# Characterization of dissolved organic matter from temperate wetlands: field dynamics and photoreactivity changes driven by natural inputs and diagenesis along the hydroperiod

**DOI:** 10.1101/2021.10.25.465760

**Authors:** Patricia E. García, Carolina F. Mansilla Ferro, María C. Diéguez

**Affiliations:** Grupo de Ecología de Sistemas Acuáticos a escala de Paisaje (GESAP) INIBIOMA, Universidad Nacional del Comahue, CONICET, Quintral 1250, San Carlos de Bariloche (8400), Argentina

**Keywords:** Patagonia, temperate wetlands, hydroperiod, dissolved organic carbon, degradation, photoreactivity

## Abstract

Wetlands store large amounts of C in biomass, sediments and water. A major C fraction is in the dissolved organic matter (DOM) pool and has multiple regulatory functions in the ecosystem. Patagonian wetlands undergo profound changes in their water cycle due to warming and reduced precipitation, causing shorter hydroperiods and reduced landscape connectivity and overall affecting their C budgets. In this study we characterized the DOM pool of a temporary wetland of North Patagonia during a hydroperiod, using optical DOM proxies obtained by absorption and fluorescence spectroscopy. DOM components were modeled through EEM-PARAFAC. DOC varied between ∼4 and ∼9 mg L^-1^, displaying aromatic signals and terrestrial/sediment fingerprints. The humic components C1 (microbial and/or vegetation derived) and C2 (soil/sediment derived) prevailed in the DOM pool, whereas the non-humic component C3 (derived from aquatic production) showed lower contribution. Along the hydroperiod DOM optical proxies allowed identifying allochthonous inputs, degradation and an increasing contribution of the internal production to the DOM pool. Photodegradation experiments showed that exposure to PAR+UVR produced slight changes in the DOC concentration and a reduction in DOM molecular weight/size. The contribution of humic *vs*. non-humic components influenced DOM photoreactivity. The prevalence of humic components determined high DOM photorecalcitrance.

## Introduction

Wetlands occur in different climates and biomes, covering around 9% of the world’s land surface, and are considered among the most threatened ecosystems by climate changes, land use, water regulation and pollution (Brinson and Malvárez 2002; Keller 2011; van Asselen *et al*. 2013; Were *et al*. 2019; Xu *et al*. 2019).

These systems experience profound seasonal variations in water level, which can involve a succession of wet and dry periods, and also undergo high inter-annual fluctuations that imply extended periods of drought or flooding. Natural climate oscillations and human-induced changes impacting on the global water cycle have profound consequences for wetlands. Temperature, evaporation, humidity, and the seasonality and amount of precipitation drive the water cycle, thereby affecting most wetland processes (van Asselen *et al*. 2013; Häder and Barnes 2019).

Globally, wetlands are estimated to store ∼35% of terrestrial carbon (C) and therefore play an important part in the global C cycle, although the pathways of C flux in these systems is still poorly understood. These ecosystems may affect the atmospheric C cycle through dynamic sequestration-emission processes (Denman *et al*. 2007). Wetlands can act as sinks for C from the terrestrial environment, can sequester CO_2_ from the atmosphere through photosynthesis of macrophytes and phytoplankton, and also transfer C to connected ecosystems. A large fraction of the total C present in wetlands is in the pool of dissolved organic matter (DOM) which includes a variety of compounds derived from the aquatic production (sediment and plant leachates, metabolites from aquatic organisms, etc.) as well as from soil and vegetation of the catchment. Macrophytes constitute a major source of particulate organic matter (POM) and DOM to the water and sediments of wetlands (Wetzel and Søndergaard 1998; Wetzel 2001; Yuan *et al*. 2020). DOM derived from aquatic production is composed of substances of comparatively lower aromaticity and molecular weight than terrigenous DOM (Cory and McKnight 2005; Stedmon and Markager 2005; Kritzberg *et al*. 2006; Murphy *et al*. 2008; Huguet *et al*. 2009). The concentration, composition and molecular structure of aquatic DOM is variable at spatial and temporal scales, and is ultimately determined by its interaction with environmental factors (climate variables, pH, conductivity, sunlight exposure), other molecules and organisms (Fellman *et al*. 2009; Chen and Jaffé 2014; Hansen *et al*. 2016). In aquatic systems the DOM pool, and particularly its light-absorptive fraction (CDOM), determines the underwater light and thermal scenarios, and thus, exerts a chief control on their heterotrophic and autotrophic production (Häder *et al*. 2011).

Temperate wetlands of the southern hemisphere have been comparatively less studied than tropical and subtropical wetlands with regard to their role in the global C cycle. In Andean Patagonia (Southern South America) temporary wetlands usually develop during autumn and last until early summer, experiencing freezing conditions at the beginning of the hydroperiod and warming up from late spring. Due to their geographical location and shallowness, their water columns are highly exposed to elevated ultraviolet (UV) radiation levels (Villafañe *et al*. 2001; Perotti *et al*. 2005; Díaz *et al*. 2006; Pérez *et al*. 2010). Challenging radiation levels enhance photochemical weathering affecting the C pool of aquatic systems, with profound effects on abiotic and biotic compartments (Zagarese *et al*. 2001; 2017; Pérez *et al*. 2003; Bastidas *et al*. 2009, 2010; Gerea *et al*. 2016).

In this investigation we focus on DOM dynamics in a cold temperate ephemeral wetland of Northwestern Patagonia over the course of a hydroperiod, aiming to characterize the natural DOM pool in terms of concentration and quality. We applied optical analysis of the light-absorbing properties of the chromophoric and fluorescent DOM fractions (CDOM and FDOM, respectively) in order to characterize the DOM quality in terms of potential sources, molecular features and diagenetic state. Additionally, we evaluated through laboratory assays the photoreactivity of natural DOM pools collected at different moments of the hydroperiod by exposing natural DOM to photosynthetically active radiation and ultraviolet radiation (PAR+UVR). We hypothesize that changes in DOM optical properties and photoreactivity along the hydroperiod reflect DOM inputs from different sources and degradation processes.

## Methods

### Study area and field sampling

The study was carried out in Fantasma Pond (FP; 41°07’ S, 71°27’ W; 798 m a.s.l.), a shallow (∼2 m), temporary piedmont wetland of Andean Patagonia (Nahuel Huapi National Park, Argentina). The climate of the area is cold temperate and markedly seasonal with ∼60% of the precipitation concentrated from May to September (hereinafter the wet season). The wet season is characterized by cold temperatures and a short photoperiod, while the dry season (spring-summer) has much lower rainfall levels, warmer temperatures and a long photoperiod with enhanced ultraviolet radiation (UVR) levels, particularly in spring (Paruelo *et al*. 1998; Villafañe *et al*. 2001).

In order to evaluate changes in the concentration and quality of the DOM pool of FP, water samples were collected monthly throughout the entire hydroperiod, from June to December. At each sampling occasion water level was recorded at the sampling point. Water temperature, pH and conductivity were measured using multiparameter probes. In November, the vertical profile of downward irradiance (380-750 nm, every 1 nm) was measured at noon with a radiometer (USB2000, Ocean Optics).

Sampling was conducted at a central point of the wetland using a Kemmerer bottle (4.7 L). Water samples were poured into acid-cleaned polycarbonate carboys which were immediately transported to the laboratory thermally insulated and in darkness. In the laboratory the samples were filtered through 0.22 µm polyvinylidene difluoride membranes (PVDF, Millipore) and stored in sealed dark flasks at 5°C for DOM characterization and photodegradation assays.

Precipitation and air temperature data for the studied period were retrieved from the meteorological station EMMA (INIBIOMA-CONICET) (Fig. S1).

### DOM photodegradation experiments

Laboratory incubations were conducted to analyze the natural DOM photoreactivity of water samples collected in FP in six consecutive months of the hydroperiod. The experimental design targeted the effect of exposure to photosynthetically active radiation and UV radiation (PAR+UVR) on the concentration and quality of natural DOM. The experiments consisted in natural DOM incubation under two treatments (each with 3 replicates): exposed to PAR+UVR or in darkness (control). Filter-sterilized DOM samples were placed in capped quartz test tubes (28 mL) and the control tubes were wrapped with aluminum foil to prevent exposure to light. The replicates were set to rotate in a custom-made wheel at 1 RPM in an environmental test chamber (Sanyo, MLR5) for 5 days at 20°C, with a photoperiod of 12 h PAR+UVR: 12 h darkness. Irradiation was provided through an array of 2 PAR lamps (Sanyo 40 W PAR = 69.1 µEm^-2^ s^-1^), 2 UV-B lamps (Q-panel 320, UVB = 4.1 kJ m^-2^) and 3 UV-A lamps (Q-panel 340 UVA = 44 kJ m^-2^) (Fig. S2a).

### Laboratory analysis

DOM characterization in terms of concentration and quality was performed on the filtered water samples collected throughout the hydroperiod and before and after exposure to the different treatments (PAR+UVR and darkness) of the photodegradation experiment.

The concentration of dissolved organic carbon (DOC) was measured in filtered water samples as non-purgeable organic carbon using a Shimadzu TOC-L high temperature analyzer with a high sensitivity catalyst (detection limit of 4 μg L^-1^).

Optical analyses were conducted to characterize the chromophoric and fluorescent DOM fractions (CDOM and FDOM, respectively). The absorbance spectra of filter-sterilized water samples were obtained at 1 nm intervals from 200 to 800 nm in a UV-Visible spectrophotometer (Hewlett Packard 8450), using a quartz cuvette (1 cm path length). ASTM1 grade water (Milli-Q) was used as a blank. For analysis of the FDOM fraction, the filtered, sterilized water samples were scanned in a Perkin-Elmer 55B spectrofluorometer equipped with a 150-W Xenon arc lamp and a Peltier temperature controller, using a 1 cm quartz fluorescence cell. Excitation-emission matrices (EEMs) were collected at specific excitation (240-450 nm at 5 nm intervals) and emission (300-600 nm at 0.5 nm intervals) wavelengths. The spectrofluorometer was set up with 10 nm excitation and emission slits and a scan speed of 1500 nm min^-1^. As detailed above, ASTM1 grade water was used as a blank.

### Data analysis

The averaged UV-visible absorbance between wavelengths 700 to 800 nm was subtracted from the absorbance spectra to correct for offsets due to several instrument baseline effects, following Helms et al. (2008). The absorbance data were converted to absorption coefficients as follows:

a_λ_ = Naperian coefficient (m^-1^) = 2.303 A_λ_/l

A = absorbance value

l = quartz cuvette path length (m) = 0.1 m

The absorption coefficient at 350 nm (a_350_) was associated with terrigenous DOM and/or with the concentration of lignin phenols (Hernes and Benner 2003; Spencer et al. 2008). The specific absorption coefficient at 350 nm (a_350_:DOC) was calculated and used as a surrogate for DOM aromatic content (Weishaar *et al*. 2003; Hansen *et al*. 2016). The spectral slopes for the intervals 275-295 nm (S_275-295_) and 350-400 nm (S_350- 400_) were calculated by fitting the log-transformed spectral data to a linear regression (Helms et al. 2008). These slopes and the slope ratio S_R_ (S_R_=S_275-295_:S_350-400_) are inversely related to DOM molecular size and aromaticity and are customarily used as proxies for DOM degradation (Helms *et al*. 2008; Spencer *et al*. 2010; Fichot and Benner 2012). The S_275-295_ and S_350-400_ and the S_R_ generally increase on irradiation (Helms *et al*. 2008; Hansen *et al*. 2016). The Climate Forcing Optical Index (CFOI) was calculated as a_320_:DOC divided by the spectral slope S_275-295_, and was applied as a proxy of sunlight/photochemical weathering and terrestrial DOM inputs (Williamson *et al*. 2014).

Two fluorescent DOM compositional indicators were calculated from the fluorescence scans of the samples. The humification index HIX, an indicator of humic content or extent of humification, was calculated from the ratio of two integrated regions of an emission scan (Em_435–480_: Em_300–345_) at Ex_254_ (Zsolnay *et al*. 1999; Ohno 2002). High HIX values indicate a high degree of humification (Hansen et al. 2016). The biological index (BIX), a proxy for autotrophic production, was calculated as the ratio of the emission intensities 380 and 430 nm (Em_380_ and Em_430_) at Ex_310_, following Huguet *et al* (2009). BIX values >1 correspond to recently-produced DOM of autochthonous origin (Hansen *et al*. 2016).

A total of 60 EEMs were processed using the software FL–WinLab^®^ (Perkin-Elmer). The absorbance spectra (200-800 nm) were used to develop a matrix of correction factors for each EEM using the Matlab FDOMcorr toolbox, accounting for inner filter effects, blank subtraction and normalization to the area under the water Raman peak of the blank at 350 nm. The resulting data were expressed in Raman units according to Murphy et al. (2010).

Parallel Factor Analysis (PARAFAC) was applied to deconvolve DOM fluorescence signals into specific components using the drEEM toolbox for MATLAB (MATLAB R2015a) (Murphy *et al*. 2013).

A principal component analysis (PCA) was used to analyze the variation in the DOM pool during the hydroperiod taking into account the water depth, the DOC concentration, CDOM parameters (a_350_:DOC, S_275-295_, S_350-400_, S_R_) and FDOM variables (HIX, BIX and the PARAFAC components). The analysis was performed using the FactoMiner package in the R environment.

A linear model was fitted to the relationship between the absorption coefficient a_350_ and DOC concentration. The results of the photodegradation assays were analyzed through two-way ANOVA, using as factors the radiation treatments (PAR+UVR and Dark) and the hydroperiod month (6 levels), using the free software R (package rstatix).

## Results and discussion

### Dynamics of the dissolved organic matter pool during the hydroperiod

The hydroperiod studied in FP lasted from June to December and reflected the precipitation and temperature patterns, reaching its maximum volume in early spring with the snowmelt water input, and then contracting until it completely dried up in early summer (Fig. S1; Table 1). Similar hydroperiod patterns have been observed in temporary wetlands adjacent to FP, which are typical of piedmont systems of the Andean Patagonian region, showing slight differences due to local fluctuations in precipitation volume (Cuassolo *et al*. 2012, 2015, 2020; Jara 2019).

The vegetation present in FP was characterized by a profuse ring of *Carex aemathorrhyncha* covering ∼60% of the pond and a mixed cover of *Eleocharis pachycarpa* and *Potentilla anserina* (∼10% and ∼30%, respectively) in the central sector.

The water level at the sampling point varied from ∼ 0.6 m to ∼1.3 m, showing higher depths in August and September (Fig. S1; Table S1). Conductivity ranged from 75 μS and 106.3 μS, with higher values in September and October coinciding with higher depths and water inputs due to increased runoff. The pH varied slightly around neutrality showing a mean value of 7.4 (±0.4) and similarly lower values at the beginning and at the end of the hydroperiod (∼6.5).

The DOC concentration ranged between ∼4 and ∼9 mg L^-1^, with higher values at the beginning and at the end of the hydroperiod, and lower values coinciding with the higher depths recorded (Fig. 1a; Table S1). These DOC levels are within the range of previously-reported concentrations for this wetland (Pérez *et al*. 2010; Cuassolo *et al*. 2012, 2020). The CDOM analysis resulted in high values of the coefficient a_350_, indicative of terrigenous DOM and/or dissolved lignin content, particularly in the samples from the beginning and the end of the hydroperiod (Fig. 1b; Table S1). These high values of a_350_ likely reflected the contribution of DOM from surrounding soils [soil organic matter (SOM)], hygrophilous plants and bottom sediments, as indicated also by the direct relationship found between DOC concentration and a_350_ (R^2^=0.98; p=0.001). The a_350_:DOC values indicated high aromaticity of the DOM pool and was sensitive to eventual external inputs, such as recorded in September following a heavy rainfall that increased the runoff (Table S1). In ephemeral wetlands such as FP it is difficult to establish what is allochthonous and autochthonous, since the system alternates between dry (terrestrial phase) and flooded periods. The major sources of POM and DOM are the soils, bottom sediments and hygrophilous plants that can tolerate drought or flooding. Within FP, *C. aemathorrhyncha* and *E. pachycarpa* tolerate the drought period and develop large stands during the flooded period. Whereas, *P. anserina* dominates during the dry phase and decays during the hydroperiod, contributing substantially to the pools of POM, DOM and nutrients. This species decomposes faster than native species leaching coloured, high molecular weight DOM (Cuassolo *et al*. 2012, 2020). Thus, inputs from the decomposition of *Potentilla* and bottom sediments are likely important DOM sources during the initial flooding, along with contributions from the terrestrial vegetation and soils surrounding the pond that are mobilized by the runoff. The bottom sediments are known to contain large amounts of organic matter (Chimner *et al*. 2011), and the leaching that occurs on inundation sends an initial pulse of DOC into the water column of wetlands. Accordingly, the initial DOM pool of FP showed optical signatures indicative of high humic content and prevalence of compounds of high molecular weight/size and aromaticity (Table S1), resembling terrigenous inputs (García *et al*. 2018).

**Figure 1.**
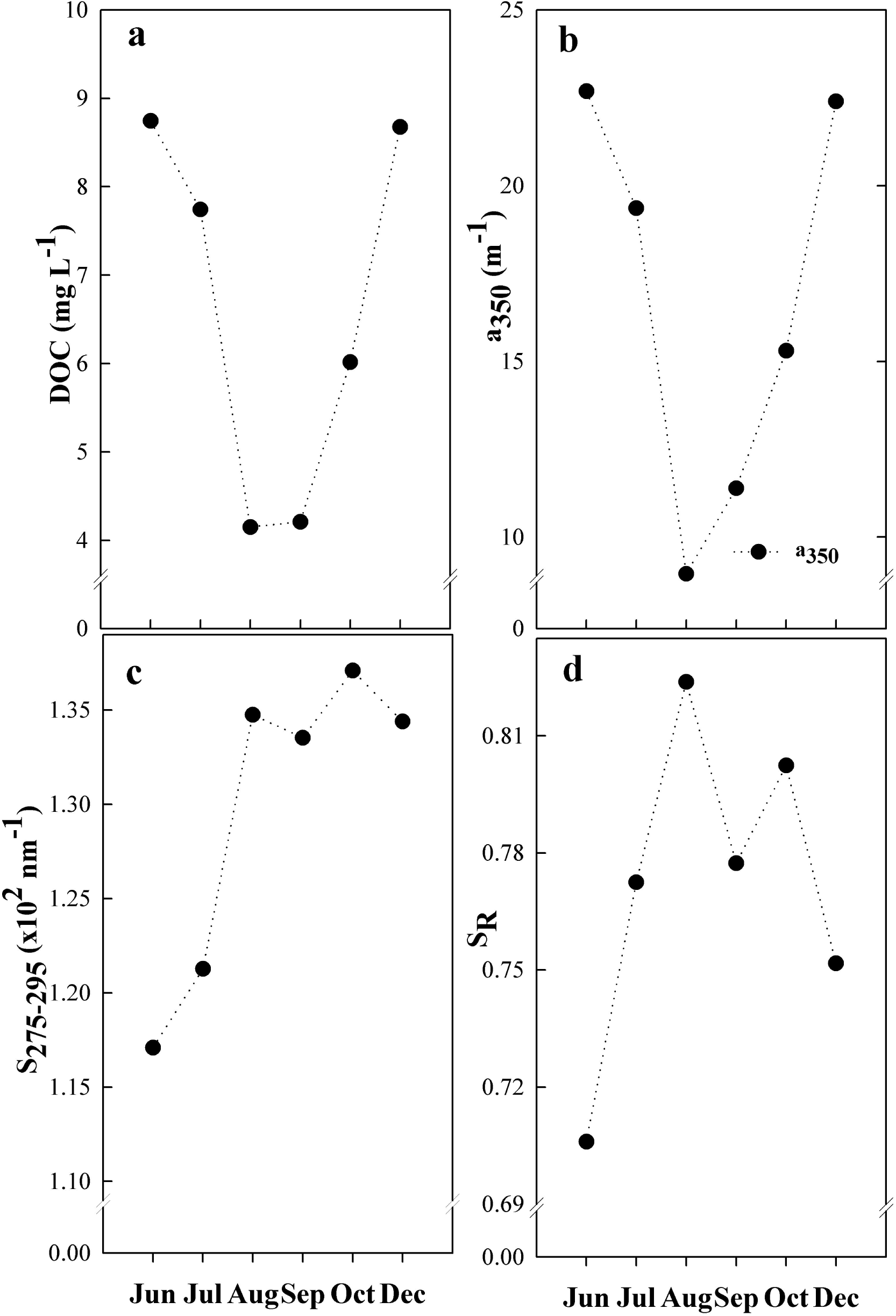
Monthly variation in different parameters of the dissolved organic matter (DOM) pool during a hydroperiod of Fantasma pond (North Patagonia, Argentina): **a)** Dissolved organic carbon (DOC) concentration; **b**) Absorption coefficients a_254_ and a_350_ (applied as proxies for aromaticity); **c)** Spectral Slope S_275-295_, and **d)** Slope ratio S_R_ (S_275-295_: S_350-400_) (applied as proxies for molecular weight and degradation).

The different optical analyses performed on the natural water samples revealed changes in DOM quality over the course of the hydroperiod. Analysis of the CDOM fraction, and particularly the increase found in the spectral slopes S_275-295,_ S_350-400_ and the S_R,_ indicated a decrease in the molecular weight/size of the DOM pool during the hydroperiod (Fig. 1c,d; Table S1). These changes point to degradation, probably the combined effect of biological and photochemical processing in the water column (Osburn *et al*. 2011; Lee *et al*. 2018). Moreover, the CFOI index, tracked the allochthonous inputs (reflected in high CFOI values) from June to August and indicated the impact of photodegradation on the DOM pool from September towards the end of the hydroperiod (lower CFOI Values). Biodegradation may be considered slow during the initial part of the hydroperiod (June to early September) due to the prevailing low temperatures (< 8°C) and even freezing conditions; however, during spring the system starts warming up, which favors biological activity, contributing to explain the stronger degradation signals observed towards the end of the hydroperiod.

In aquatic systems of Andean Patagonia, UV radiation is a limiting factor for aquatic biota and a key driver of biochemical processes in the water column, constituting a governing factor of DOM transformation (Pérez *et al*. 2003; Marinone *et al*. 2006; Zagarese *et al*. 1998, 2017). The region experiences high exposure to solar radiation in spring and summer and high UV levels year-round due to its lati/altitudinal location (Villafañe *et al*. 2001; Díaz *et al*. 2006). Moreover, from spring to autumn the longer photoperiod with high UV levels enhances photochemical processes in the water column (Queimaliños *et al*. 2019). In shallow wetlands a major fraction of the UV radiation is absorbed in the first 10-20 cm (Fig. S2a), however, the water column is mixed by the action of strong winds, which enhances photochemical weathering (Zagarese *et al*. 1998, 2017). In spring, the pond displays a lower DOC concentration due to the increase in water input as a consequence of snowmelt, promoting diluted conditions and enhancing light penetration in the water column; such a pattern has been observed in shallow lakes of the area (Soto Cárdenas *et al*. 2017).

The analysis of the FDOM fraction also supported the observed changes in DOM quality reported above. First, the outcome of the PARAFAC modelling showed that the FDOM pool of FP is composed of two humic-like components (C1 and C2) and one non-humic component (C3) (Table S2). The fluorescent components C1 and C2 showed two excitation maxima at a single emission spectrum, as a combination of two fluorescent peaks [C1 (A + M peaks) and C2 (A + C peaks)]. Whereas, C3 was characterized by a single peak, corresponding to peak T. The fluorescent peaks A and M included in C1 likely correspond to humic substances originating from microbial activity, while peaks A + C included in C2 have been attributed to terrestrially-derived humic compounds. Component C3, characterized by the T peak, has been associated with non-humic compounds of autochthonous origin (aquatic production) (Table S2). Throughout the hydroperiod the humic components C1 and C2 prevailed in the DOM pool, whereas C3 showed a smaller contribution (Fig. 2b).

**Figure 2.**
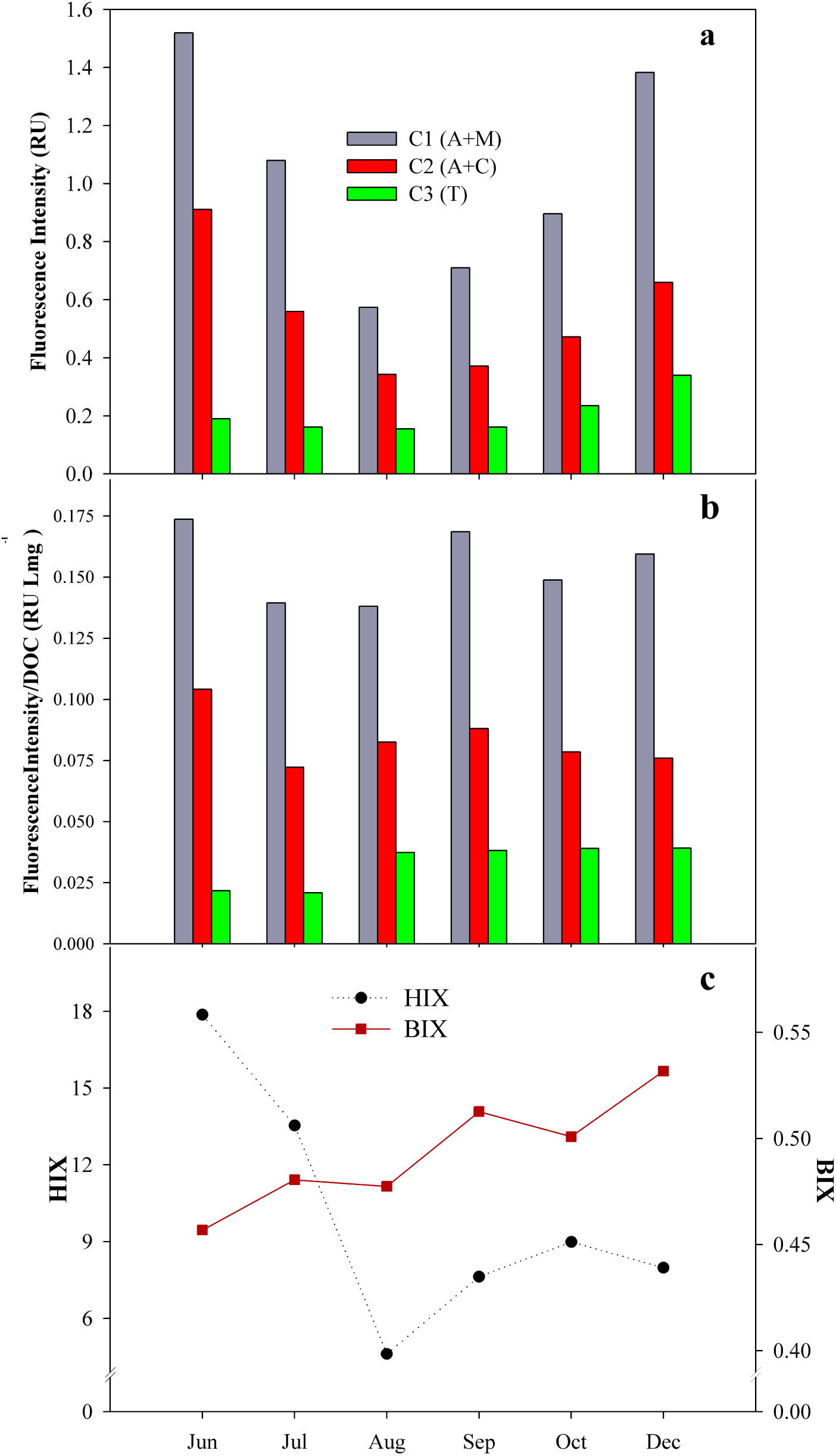
Variation in FDOM parameters over a hydroperiod of Fantasma pond: **a**) Intensity of the PARAFAC components C1, C2 and C3; **b**) DOC-normalized intensity of the FDOM components C1, C2 and C3, and **c**) Humification index (HIX: black dotted line) and Biological Index (BIX: red solid line).

The intensities of the three fluorescent components followed the DOC concentration pattern (Fig. 1a; Fig. 2a). However, the DOC-normalized components displayed a relatively stable and prevalent contribution of the humic components C1 and C2 and an increase in C3 during the hydroperiod (Fig. 2b). This pattern was supported by the high HIX values highlighting the contribution of vegetation and soil/sediment DOM. In addition, the increasing contribution of C3 and BIX mirrored the growing input of the aquatic production over the course of the hydroperiod (Fig. 2b, c).

The PCA analysis performed to analyze variation in the DOM pool during the hydroperiod, taking into account DOC concentration, CDOM parameters and FDOM variables, resulted in two valid principal components (PC). PC1 and PC2 contributed to explain 57.2% and 29.9%, respectively, accounting to explain 87.11% of the total variance (Table S3). PC1 correlated positively with the DOC concentration, a_350_:DOC, C1, C2, and HIX, and showed a negative correlation with water depth, S_275-295_ and S_R_ (Fig. 3; Table S3). PC2 correlated positively with the degradation proxies S_275-295_ and S_350-400_ and with the proxies of autochthonous production, C3 and BIX. Along the PC1 axis two groups of samples were distinguished. On the right of the plot, samples from June, July and December were grouped, based on their higher DOC, a_350_:DOC, high C1 and C2 intensities and HIX and corresponding with lower depths of the wetland. Based on contrasting values of the contributing variables, samples from August, September and October were clustered on the left of the plot (Fig. 3). PC2 separated the samples from September, October and December with higher values of S_350-400_, C3 and BIX, indicative of higher degradation and the contribution of autochthonous biological activity (Fig. 3). Overall, the PCA allowed discriminating samples with high DOC concentrations supported by differences in DOM quality. Samples from the beginning of the hydroperiod, derived from vegetation and soil/sediment inputs, showed a strong humic fingerprint. In contrast, those from the end of the hydroperiod reflected autochthonous production. Regarding the vegetation inputs, it is difficult to discriminate between the contributions of terrestrial and hygrophilous vegetation to the DOM pool of the pond, since they are both sources of high molecular weight and aromatic DOM due to their lignin and cellulose contents (Talbot and Treseder 2012; Hansen *et al*. 2016). Collectively, these results indicate that over the course of the hydroperiod, the DOM pool of FP showed pronounced concentration and quality changes. Although the prevailing DOM sources appear to be terrestrial and hygrophilous vegetation and the soils and bottom sediments, the contribution of aquatic production increases from early spring towards the end of the hydroperiod. The aquatic contribution could be tracked through the increase in the C3 and BIX, likely enhanced by the activity of aquatic vegetation, microbiota (plankton, sediment microbiota, epiphytic microbial communities, etc.) and their metabolites. The changes observed in the concentration and quality of the DOM pool along the hydroperiod suggested that inputs and degradation might trigger DOM reactivity changes. This hypothesis was tested through photodegradation experiments performed with DOM samples collected monthly during the hydroperiod.

**Figure 3.**
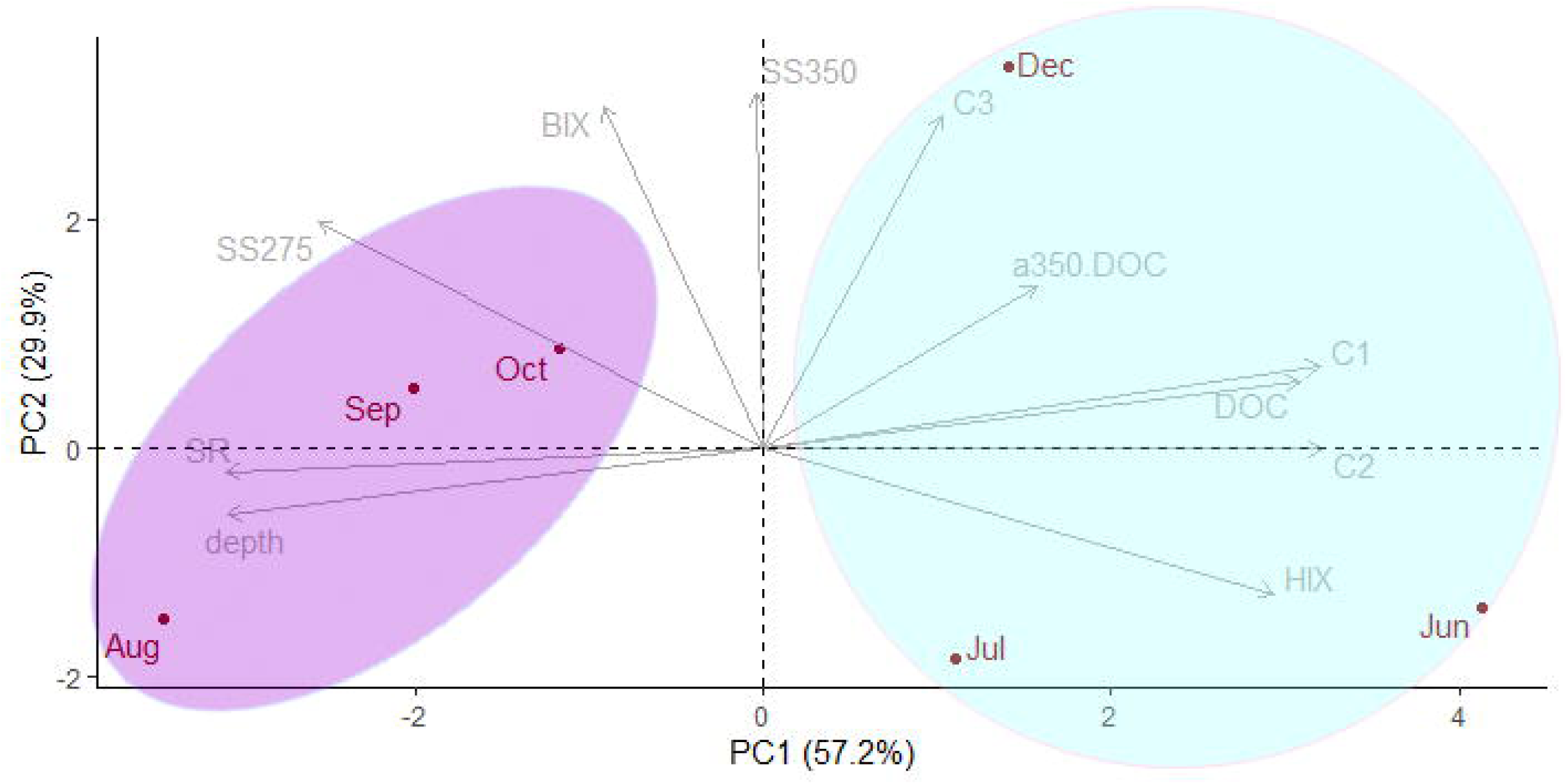
Plot of the Principal Component Analysis (PCA) PC1 *vs*. PC2 performed to study variation in the DOM pool of Fantasma Pond over a hydroperiod, including DOC concentration and CDOM (a_254_; a_350_; SUVA; a_350_:DOC; S_275-295_; S_350-400_ and S_R_) and FDOM proxies (fluorescent components C1, C2 and C3; and the humification and biological indexes: HIX and BIX).

### Photodegradation experiments

Photodegradation experiments were performed by exposing water samples to PAR+UVR and dark treatments. Significant differences were detected in CDOM and FDOM proxies between treatments and across samples from different months of the hydroperiod (Table S4), suggesting that natural changes in DOM concentration and quality influenced the magnitude of photodegradation.

The effect of irradiation on DOC concentration was rather idiosyncratic among the water samples studied. Notably, samples from September and December showed a significant reduction in DOC concentration on exposure to PAR+UV, whereas in the other samples the trend was opposite or not significant (Fig. 4a).

**Figure 4.**
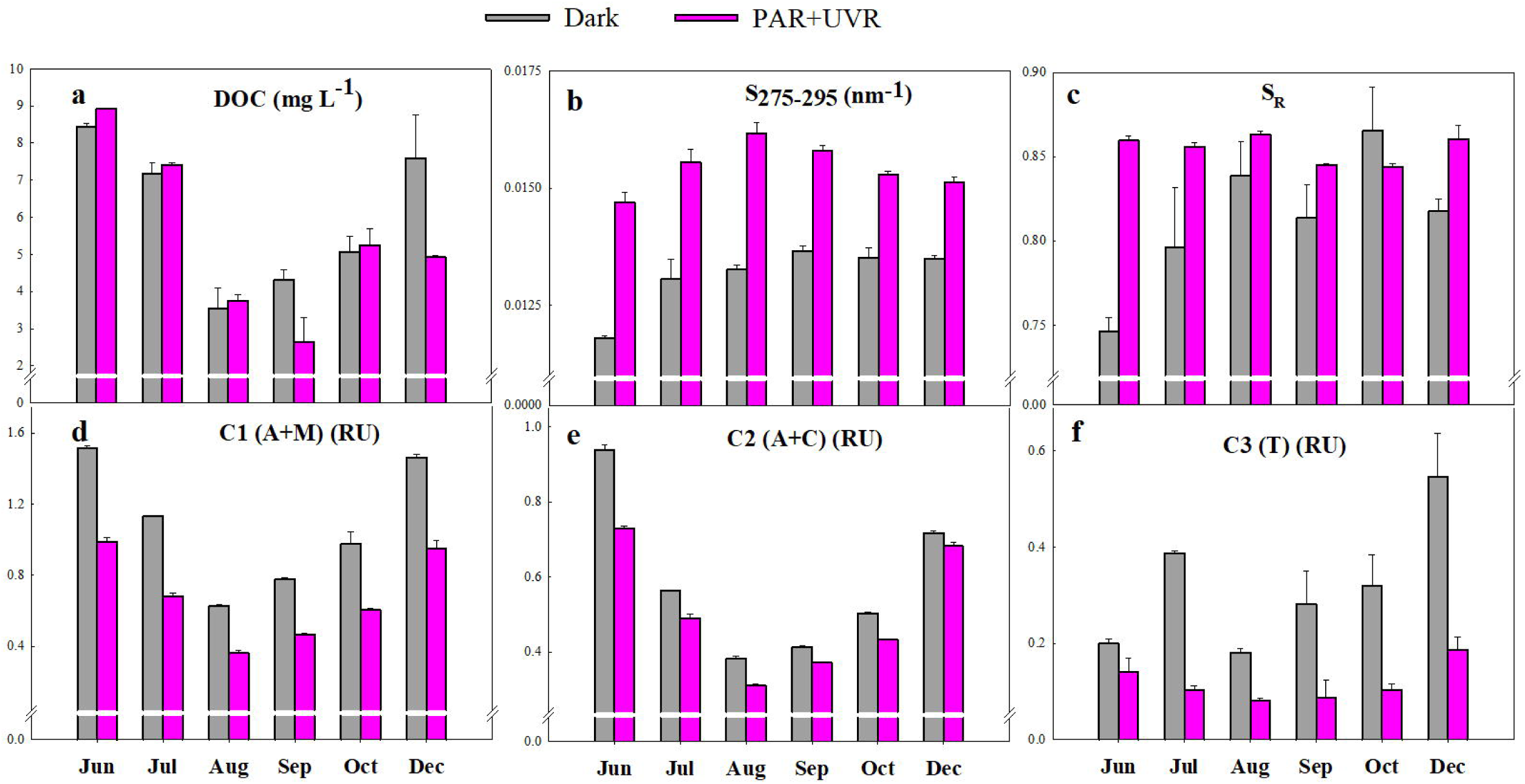
Changes in DOM properties in laboratory incubations under PAR+UVR and DARK treatments performed to study the DOM photoreactivity of DOM samples collected monthly during a hydroperiod in Fantasma Pond: **a**) DOC concentration**; b**) Spectral slope S_275-295_; **c**) Slope ratio S_R_ (S_275-295_: S_350-400_); **d**) fluorescent component C1; **e**) fluorescent component C2, **f**) fluorescent component C3.

In terms of changes in DOM quality, the PAR+UV treatment produced a significant loss of absorbance in the water samples, reflected in lower values of the a_350_:DOC, and a significant increase in S_275-295_ and S_R,_ indicative of a reduction in DOM molecular weight/size (Table S4; Fig.4 b, c). The intensity of the three fluorescent components decreased significantly on irradiation; however, differences in the loss of fluorescence were detected among the three components (Fig. 4 d, f; Table S4). The humic component C1 and the non-humic C3 showed higher loss of fluorescence intensity than the humic component C2, which was comparatively more stable or photoresistant (Fig. 4d-f).

The negative slopes of the relationship between a_350_ and S_275-295_ computed for all the incubations (Fig. 5a) indicated that exposure to radiation consistently reduced DOM molecular weight/size, although the range of the slope values suggested differences in the degree of transformation among samples, even in those with similar a_350_ values (Fig. 5a). The relationship a_350_ vs. S_275-295_ allowed the detection of differences in photo-degradation of the DOM pools from different moments of the hydroperiod, regardless of DOC concentration. The magnitude of photodegradation reflects changes in DOM, likely related to its composition and/or diagenetic state. This idea is also supported by the relationship found between the intensity of the three fluorescent components and S_275-295_, which showed differences in photoreactivity. Steeper slopes were found in the case of C1 and C3, while a smoother pattern was observed in the case of C2, indicating its photo-resistant condition (Fig. 5 b-d). The photoreactivity of the components can therefore be characterized as C1>C3>C2. The humic component C1 includes compounds probably derived from microbial sources, and was more photolabile/photoreactive than the humic component C2, which has a stronger soil/sediment fingerprint. Furthermore, the stability displayed by C2 in the DOM pool throughout the hydroperiod may be taken as an indication of its recalcitrance also to biological degradation (Fig. 2b). Other investigations focusing on the photoreactivity of DOM in running waters and shallow permanent lakes in the same catchment have also reported high recalcitrance of the humic component C2 (or A+C peaks) (Soto Cárdenas *et al*. 2017; García *et al*. 2018).

**Figure 5.**
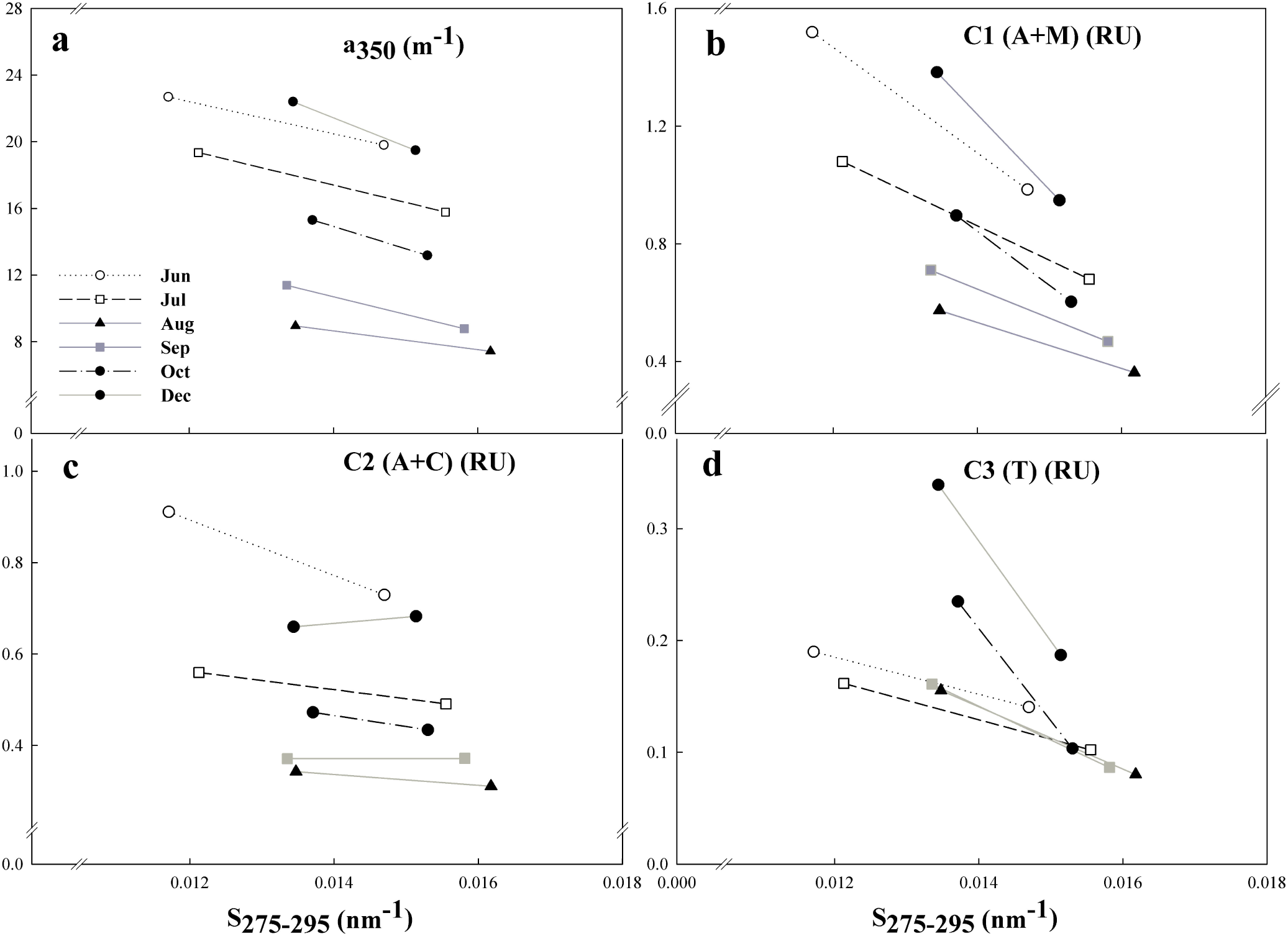
Relationship between the absorption coefficient a_350_ (**a**), the fluorescent components C1 (**b**), C2 **(c**) and C3 (**d**) and the spectral slope S_275-295_ in DOM samples collected monthly from Fantasma Pond and incubated under PAR+UVR in the laboratory.

## Conclusions

The DOC concentration and quality of the wetland varied markedly over the hydroperiod in response to inputs from terrestrial, hygrophilous vegetation and soil/sediment DOM.

Higher DOC concentrations were found at the beginning and at the end of the hydroperiod, coinciding with the lower water levels, while lower DOC concentrations were recorded at the end of winter and in early spring, following the rise in water level due to snowmelt.

The DOM pool of the wetland was characterized by the prevalence of high molecular weight, aromatic DOM with strong soil/sediment and vegetation (terrestrial and hygrophilous) fingerprints.

The progression of degradation and a low but increasing contribution of internal production to the DOM pool was detected over the hydroperiod.

In laboratory experiments, DOM photochemical transformation involved only minor changes in DOC concentration but significant changes in quality, as indicated by several optical proxies of degradation and biological production.

Changes in the relative contribution of the humic *vs*. non-humic components, specifically by the components C1 and C3, determined the reactivity and photo-degradability (C1>C3>C2) of the DOM pools at different moments of the hydroperiod.

Our results suggest that the cold temperate wetlands of Andean Patagonia can process a moderate amount of the C pool they receive/produce due to the combined effect of their DOM quality (highly recalcitrant) and the low temperature that prevails during most of the hydroperiod, which dampens biodegradation. Despite the importance of photodegradation processes, the high resistant DOM pool prevents photo-mineralization during the short hydroperiod, leading to accumulation of recalcitrant material in the sediments that may be subjected to weathering and microbial processing during the dry phase.

## Acknowledgments

We thank the town council of San Carlos de Bariloche for granting permission to collect samples in Fantasma pond, and to Mrs Audrey Shaw for language revision of the manuscript.

