## Supplementary material for "Characterization of dissolved organic matter from temperate wetlands: field dynamics and photoreactivity changes driven by natural inputs and diagenesis along the hydroperiod": Fig. S

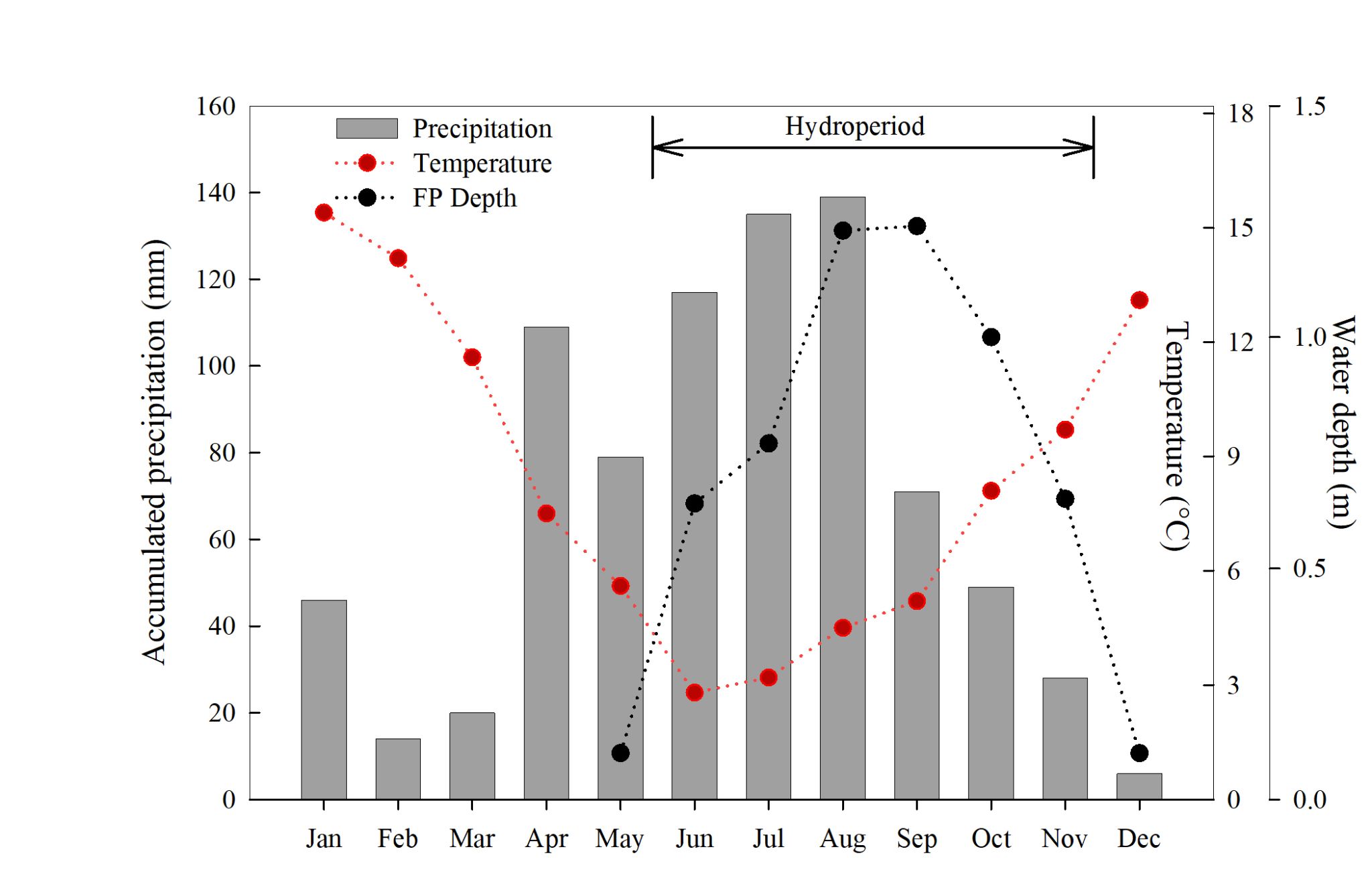

**Figure S1.** Monthly accumulated precipitation (gray bars), mean temperature (red circles with line) and water depth (black circles with dotted line) during 2014.

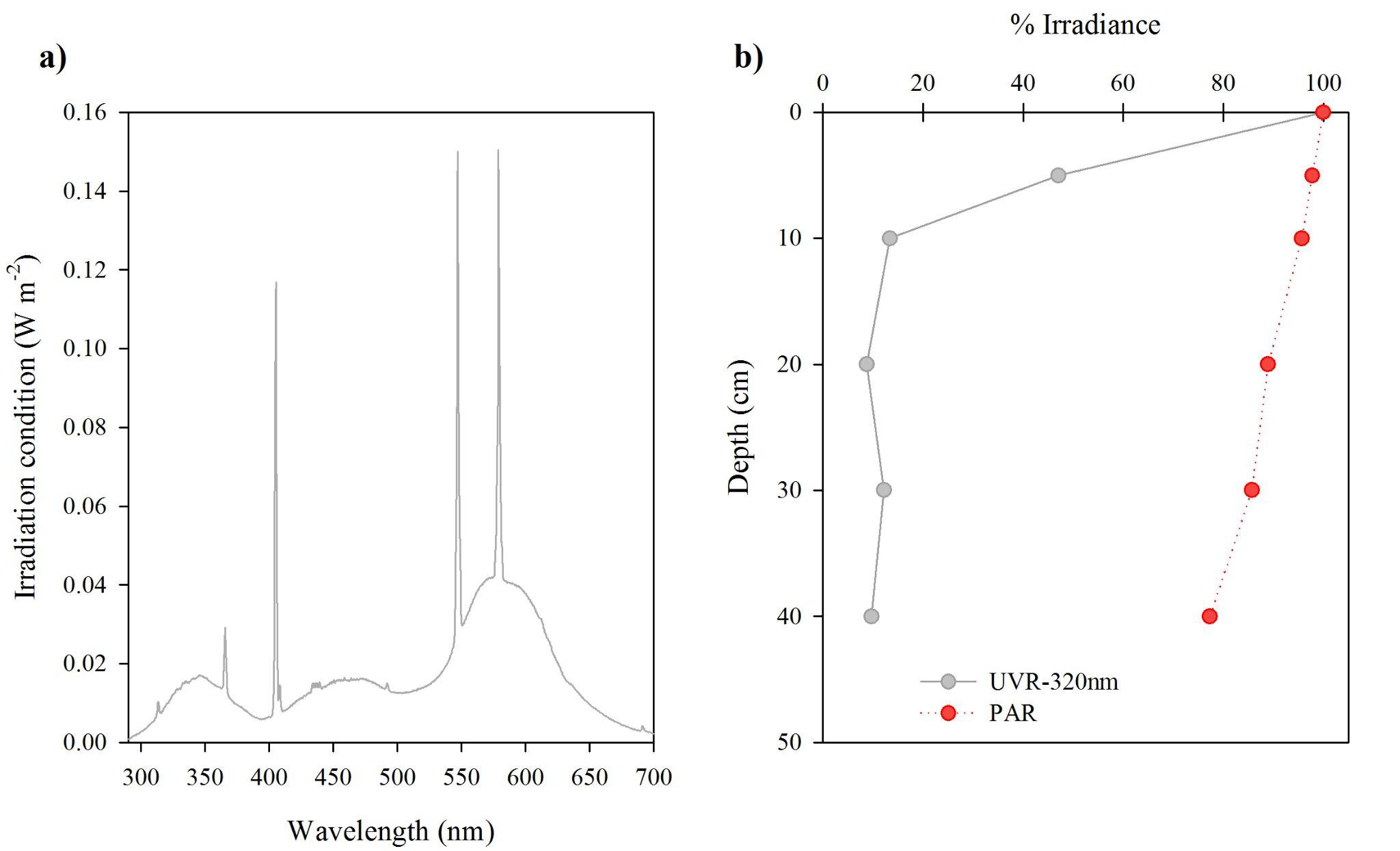

**Figure S2.** a) Radiation conditions applied in laboratory experiments; b) Vertical profile of photosynthetically active radiation (PAR) and ultraviolet radiation at 320 nm (%) in a spring sampling date (November) in the studied wetland.

**Table S1.** Monthly Variation of different DOM parameters along the hydroperiod of Laguna Fantasma (North Patagonia, Argentina). DOC: dissolved organic carbon concentration; a_350_: absorption coefficient at 350 nm; a_350_:DOC (DOC-normalized absorption coefficient at 350 nm); S_275-295_: spectral slope between 275 and 295 nm; S_350-400_: spectral slope between 350 and 400 nm; SR (slope ratio; S_275-295_: S_350-400_); CFOI (Climate Forcing Optical Index); fluorescent components C1, C2 and C3; HIX (humification index); BIX (biological index) and maximum depth of the water column (m).

|  | **a350**  **(m^-1^)** | **DOC**  **(mg L^-1^)** | **a350:DOC** | **S_275-295_**  **(nm^-1^)** | **S_350-400_**  **(nm^-1^)** | **SR** | **CFOI**  **nm m^2^ (g C)^-1^** | **C1(A+M)**  **(R.U.)** | **C2(A+C)**  **(R.U.)** | **C3(T)**  **(R.U.)** | **HIX** | **BIX** | **Depth (m)** |
| --- | --- | --- | --- | --- | --- | --- | --- | --- | --- | --- | --- | --- | --- |
| **June** | 22.68 | 8.75 | 2.593 | 0.0117 | 0.017 | 0.71 | 172.85 | 1.52 | 0.91 | 0.19 | 17.87 | 0.457 | 0.64 |
| **July** | 19.35 | 7.74 | 2.500 | 0.0121 | 0.016 | 0.77 | 178.72 | 1.08 | 0.56 | 0.16 | 13.54 | 0.481 | 0.77 |
| **August** | 8.95 | 4.15 | 2.157 | 0.0135 | 0.016 | 0.82 | 248.27 | 0.57 | 0.34 | 0.16 | 4.61 | 0.477 | 1.23 |
| **September** | 11.39 | 4.21 | 2.705 | 0.0134 | 0.017 | 0.78 | 78.76 | 0.71 | 0.37 | 0.16 | 7.62 | 0.513 | 1.24 |
| **October** | 15.30 | 6.02 | 2.544 | 0.0137 | 0.017 | 0.80 | 142.85 | 0.90 | 0.47 | 0.23 | 8.98 | 0.501 | 1 |
| **December** | 22.39 | 8.68 | 2.581 | 0.0134 | 0.018 | 0.75 | 74.18 | 1.38 | 0.66 | 0.34 | 7.97 | 0.532 | 0.65 |

**Table S2.** Description of the three fluorescent components modelled through Parallel Factor Analysis from EEMs obtained from natural DOM samples collected monthly in Fantasma Pond along a hydroperiod. C1: fluorescent component 1; C2: fluorescent component 2; C3: fluorescent component 3.

|  | **Ex_max_**  **(nm)** | **Em_max_**  **(nm)** | **Loading** | **Referenced peaks name and maxima range** | **Description** |
| --- | --- | --- | --- | --- | --- |
| **C1** | 310 (240) | 418.5 | 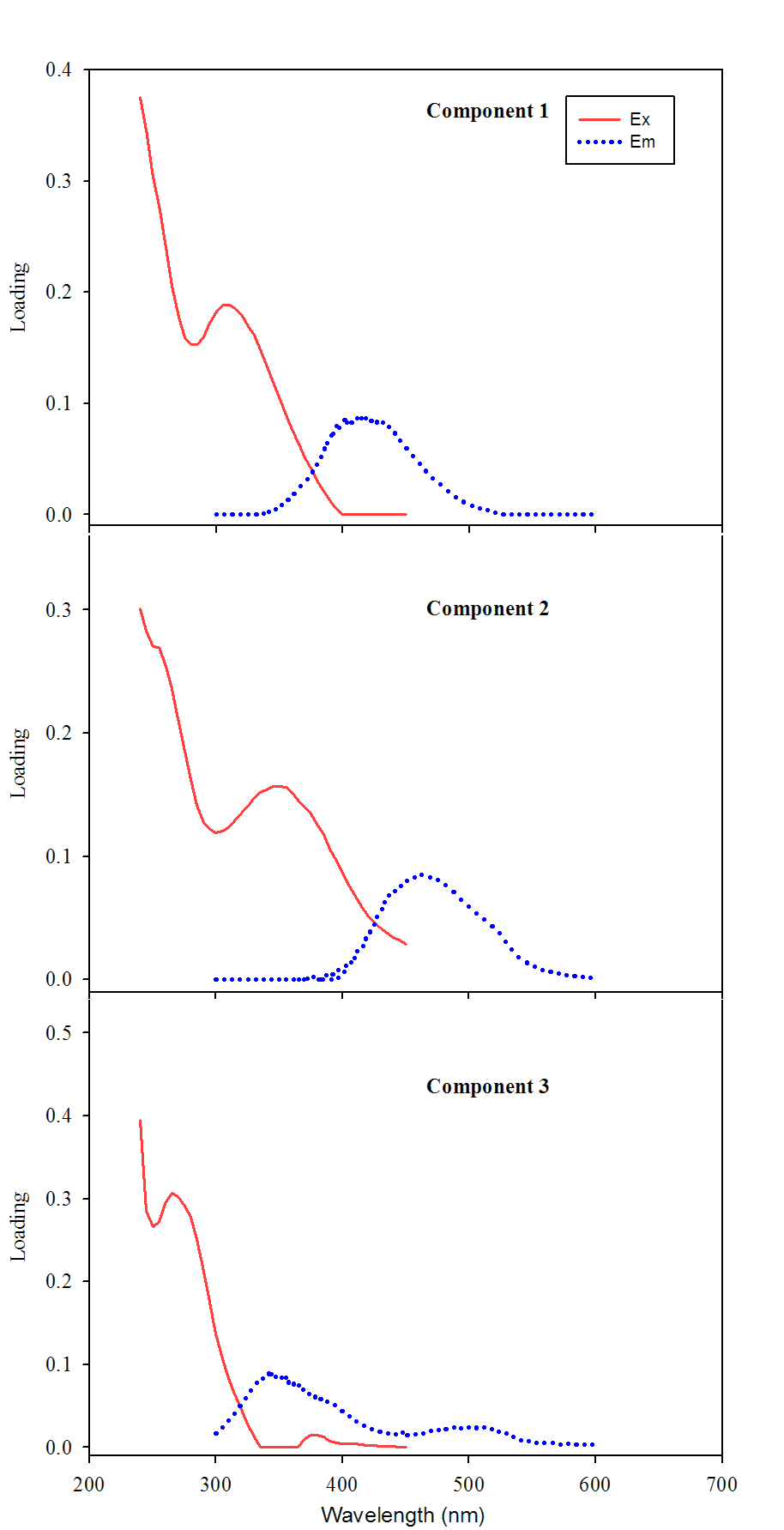 | **M peak** Ex_max_=290-310;  Em_max_= 370-420  **A peak** Ex_max_=230-260;  Em_max_=380-465 | Humic-like/ biological degradation |
| **C2** | 355 (240) | 462 | 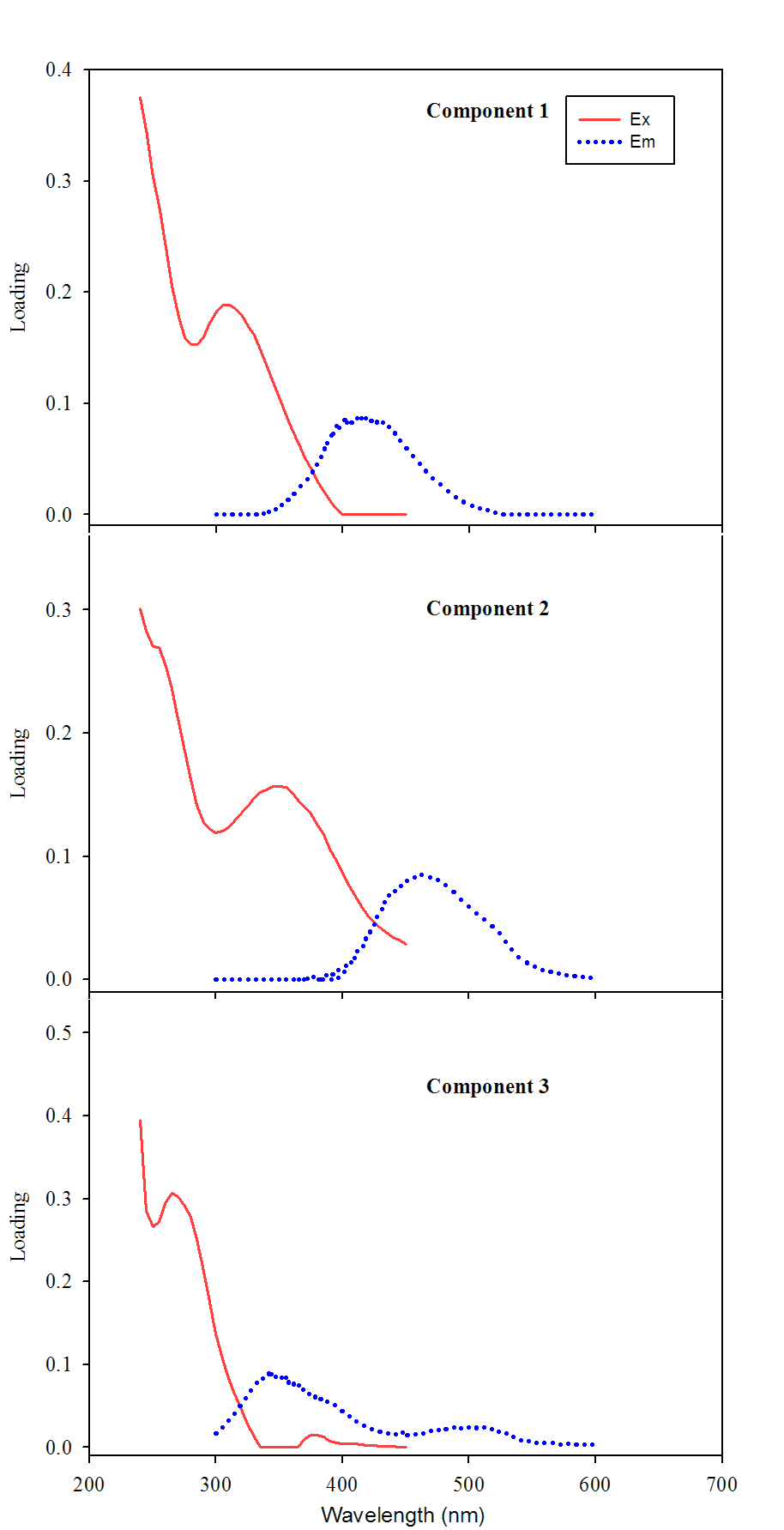 | **C peak** Ex_max_=320-360;  Em_max_ 420-480  **A peak** Ex_max_=230-260;  Em_max_=380-465 | Terrestrial humic-like |
| **C3** | 275 | 349.5 | 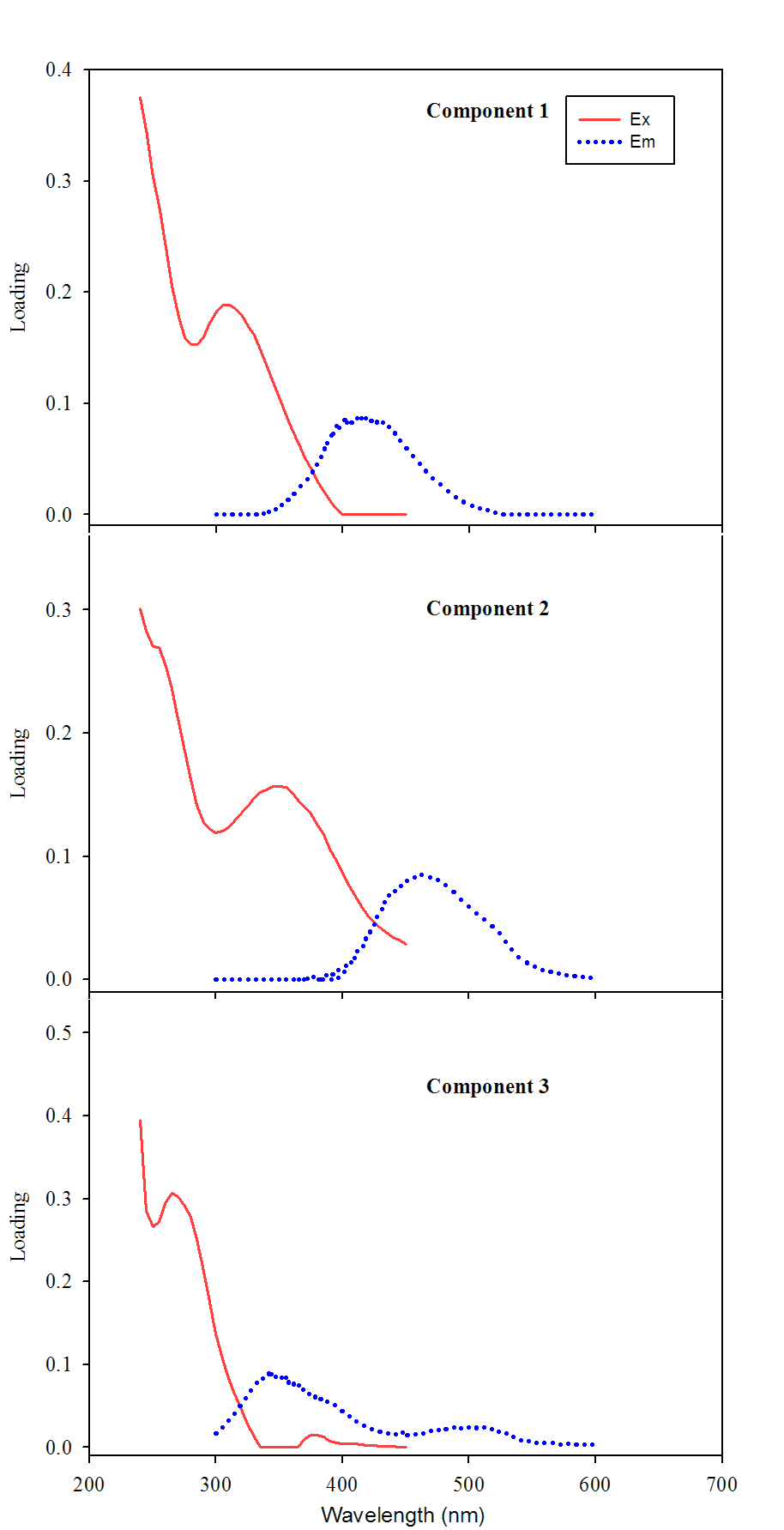 | **T peak** Ex_max_=270-280;  Em_max_=330-368 | Protein-like (non-humic) |

**Table S3:** Results of the Principal Component Analysis (PCA) performed to study the variation of optical features of the natural DOM pool in Fantasma Pond along a hydroperiod. References: DOC: dissolved organic carbon concentration; a_350_: absorption coefficient at 350 nm; S_275-295_: spectral slope for the interval 275-295 nm; S_350-400_: spectral slope for the interval 350-400 nm; SR: slopes ratio; C1, C2 and C3: PARAFAC components; BIX: Freshness index; HIX: Humification index; Depth: water column maximum depth.

| **Variables** | **PC 1** | **PC 2** |
| --- | --- | --- |
| C2(A+C) | **0.97** | -0.01 |
| C1(A+M) | **0.96** | 0.21 |
| DOC | **0.93** | 0.17 |
| C3(T) | 0.31 | **0.88** |
| HIX | **0.88** | -0.38 |
| Depth | **-0.93** | -0.17 |
| S_R_ | **-0.93** | -0.06 |
| BIX | -0.28 | **0.90** |
| S_275-295_ | -0.77 | 0.59 |
| a_350_:DOC | 0.47 | 0.43 |
| S_350-400_ | -0.01 | **0.93** |
| Eigenvalue | 6.29 | 3.28 |
| Variance explained (%) | 57.2 | 29.90 |
| Cumulative Variance explained (%) | **57.2** | **87.11** |

**Table S4.** Results of the Two-way ANOVA performed to study the changes in optical DOM parameters in samples collected monthly in Fantasma Pond during a hydroperiod and exposed to PAR+UVR and Dark treatments in laboratory incubations.

| **Parameter** | **Factor** | **DF** | **F** | **P-value** |
| --- | --- | --- | --- | --- |
| **DOC** | Treatment | 1 | 11.39 | **0.003** |
|  | Month | 5 | 113.12 | **<0.001** |
|  | Treatment*Month | 5 | 11.29 | **<0.001** |
| **a_350_** | Treatment | 1 | 956.84 | **<0.001** |
|  | Month | 5 | 1892.09 | **<0.001** |
|  | Treatment* Month | 5 | 16.19 | **<0.001** |
| **s_275-295_** | Treatment | 1 | 1246.29 | **<0.001** |
|  | Month | 5 | 46.2 | **<0.001** |
|  | Treatment* Month | 5 | 11.66 | **<0.001** |
| **S_R_** | Treatment | 1 | 496.02 | **<0.001** |
|  | Month | 5 | 17.84 | **<0.001** |
|  | Treatment* Month | 5 | 36.89 | **<0.001** |
| **a_350_:DOC** | Treatment | 1 | 2.77 | **0.109** |
|  | Month | 5 | 11.99 | **<0.001** |
|  | Treatment* Month | 5 | 7.1 | **<0.001** |
| **C1 (A+M)** | Treatment | 1 | 1969.63 | **<0.001** |
|  | Month | 5 | 740.25 | **<0.001** |
|  | Treatment* Month | 5 | 23.83 | **<0.001** |
| **C2 (A+C)** | Treatment | 1 | 1521.5 | **<0.001** |
|  | Month | 5 | 5141.8 | **<0.001** |
|  | Treatment* Month | 5 | 148.6 | **<0.001** |
| **C3 (T)** | Treatment | 1 | 215.19 | **<0.001** |
|  | Month | 5 | 23.76 | **<0.001** |
|  | Treatment* Month | 5 | 10.93 | **<0.001** |
| **BIX** | Treatment | 1 | 421.77 | **<0.001** |
|  | Month | 5 | 42.31 | **<0.001** |
|  | Treatment* Month | 5 | 5.22 | **0.002** |
| **HIX** | Treatment | 1 | 10.88 | **0.003** |
|  | Month | 5 | 17.84 | **<0.001** |
|  | Treatment* Month | 5 | 4.274 | **0.006** |
